# ZapA tetramerization is required for midcell localization and ZapB interaction in *Escherichia coli*

**DOI:** 10.1101/749176

**Authors:** Nils Y. Meiresonne, Tanneke den Blaauwen

**Author notes:** Bacterial Cell Biology, Instituto de Tecnologia Química e Biológica António Xavier, Universidade nova de Lisboa, Oeiras, Portugal (Nils Y. Meiresonne).

## Abstract

Bacterial cell division is guided by FtsZ treadmilling precisely at midcell. FtsZ itself is regulated by FtsZ associated proteins (Zaps) that couple it to different cellular processes. ZapA is known to enhance FtsZ bundling but also forms the synchronizing link with chromosome segregation through ZapB and *matS* bound MatP. ZapA exists as dimers and tetramers in the cell. Using the ZapA^I83E^ mutant that only forms dimers, this paper investigates the effects of ZapA multimerization state on its interaction partners and cell division. By employing (fluorescence) microscopy and Förster Resonance Energy Transfer *in vivo* it is shown that; dimeric ZapA is unable to complement a *zapA* deletion strain and localizes diffusely through the cell but still interacts with FtsZ that is not part of the cell division machinery. Dimeric ZapA is unable to recruit ZapB, which localizes in its presence unipolarly in the cell. Interestingly, the localization profiles of the chromosome and unipolar ZapB anticorrelate. The work presented here confirms previously reported *in vitro* effects of ZapA multimerization *in vivo* and further places it in a broader context by revealing the strong implications for ZapB localization and *ter* linkage.

## 1. Introduction

ZapA is broadly conserved among Gram-negative and -positive bacteria [1,2]. *E. coli* ZapA promotes FtsZ polymerization through enhancing cooperativity of FtsZ polymer association [3]. However, the turnover of FtsZ is highly dynamic and its filaments would thus not benefit from being rigid when constriction occurs [4]. Many of the ZapA enhanced FtsZ polymerization studies were performed *in vitro* at non-physiological conditions that themselves allow FtsZ to filament and bundle [2]. Experiments performed under physiological conditions revealed a more dynamic stabilizing effect of ZapA on FtsZ bundle formation [3].

An outstanding question is whether the proposed stabilizing effects of ZapA influence FtsZ treadmilling dynamics. Very recently, *in vitro* work showed that transient interactions of ZapA with FtsZ increases the spatial order and stabilizes the architecture of the FtsZ filament network without affecting its treadmilling velocity [5]. These effects were only observed for tetrameric ZapA that was able to interact with FtsZ [5]. ZapA can exist as a mixture of dimers and tetramers *in vitro* but *in vivo* it is likely to be mostly tetrameric due to molecular crowding conditions [3,6,7]. In fact, the available ZapA structures and *in vitro* cross-linking showed a tetrameric structure and tetramerization has already been suggested to be required for FtsZ bundling [6,7,8,9]. These *in vitro* results are interesting but also prompt an *in vivo* explanation of ZapA tetramerization functionality. ZapA acts on FtsZ as a component of the chromosome terminus of replication (*ter*) linkage that synchronizes cell division with chromosome segregation [10]. Indeed, ZapA interacts with ZapB, which interacts with MatP, which binds 23 *matS* sequences distributed on the ter domain and condenses this region of the chromosome [11–13]. Therefore, ZapA multimerization dynamics should be investigated in the context of both cell division and chromosome segregation.

Here we describe the molecular behavior of ZapA and its interacting proteins *in vivo*. Dimeric ZapA was unable to complement the ZapA deletion phenotype. Although it did not localize to the division site, it was able to interact with FtsZ elsewhere in the cell, titrating some of it away from midcell. ZapB midcell localization was lost in cells with dimeric ZapA but did not show the same diffuse localization pattern as the ZapA dimers. Instead it resided predominantly at one cell pole suggesting ZapA tetramerization is important for ZapB interaction. Finally, we observed that chromosomal localization is affected in cells without ZapA or with dimeric ZapA as it anticorrelates with the polar ZapB.

## 2. Results

### 2.1. Dimeric ZapA^I83E^ does not complement the ΔzapA phenotype

The work of Pacheco-Gómez *et al* [6] shows that ZapA tetramerization is required for FtsZ bundling *in vitro* using ZapA mutant I83E that only forms dimers. This mutant fully folds and forms ZapA dimers that were shown to still bind FtsZ by co-sedimentation [6]. To assess whether ZapA^I83E^ would complement the elongated Δ*zapA* phenotype and restore wild-type morphology *in vivo*, a complementation experiment was performed. Expression of WT ZapA, ZapA^I83E^ or a negative empty vector (EV) control was induced from plasmid with 50 μM IPTG in TB28 Δ*zapA* cells growing in rich medium for ~8 mass doubling as described before [7]. The cells were then fixed, imaged, and average cell lengths were analyzed. This showed that ZapA^I83E^ was unable to complement Δ*zapA* with 6 % of the cells being longer than 10 μm compared to wild type ZapA with 1 % cells longer than 10 μm. ZapA^I83E^ results resembled more the EV control, which had 8 % of the cells longer than 10 μm and ZapA results resembled more the TB28 parental strain with 0.5 % cells longer than 10 μm (**Figure 1**).

**Figure 1.**
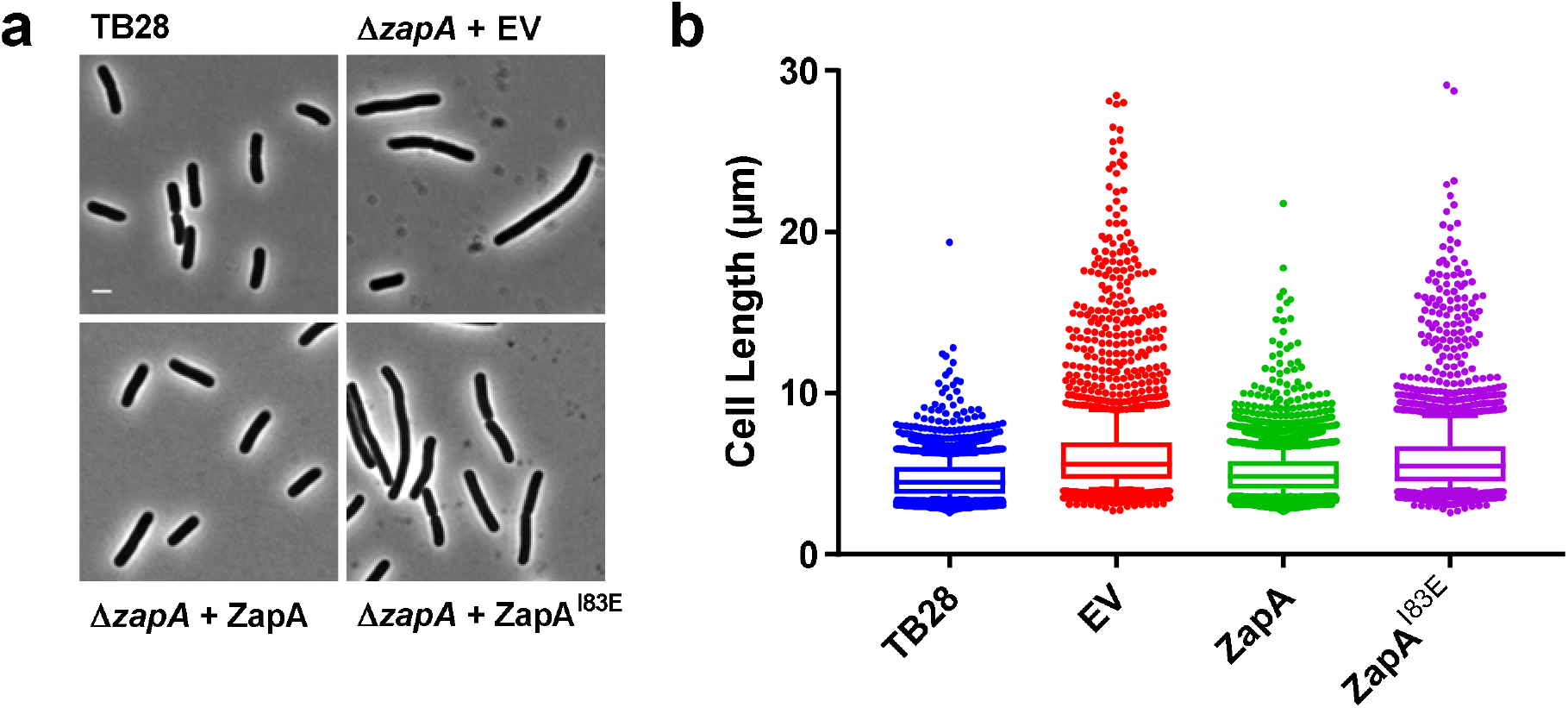
ZapA^I83E^ does not complement the elongated Δ*zapA* phenotype. **(a)** Phenotypes of the cells. The scale bar represents 2 μm. **(b)** Cell length distributions of each group. The number of cells and their average length measured were for the wild type parental strain TB28; 3670 and 4.5 μm, EV; 2686 and 6.4 μm, from plasmid expressed ZapA^WT^; 4476 and 5.1 μm and ZapA^I83E^; 2491 and 6.1 μm, respectively. The whiskers represent the 10^th^ and 90^th^ percentile.

### 2.2. Dimeric ZapA^I83E^ does not localize at midcell

ZapA^I83E^ was unable to complement a *zapA* deletion strain based on average cell length. *In vitro* work suggested that dimeric ZapA^I83E^ is still be able to bind FtsZ [6]. Therefore, it was hypothesized that ZapA^I83E^ would localize with FtsZ at midcell and that this may obstruct proper divisome functionality. The Δ*zapA* cells from the complementation experiment were immunolabeled with antibodies against FtsZ, ZapA, or ZapB and a fluorescent secondary antibody to probe their localization patterns. This revealed that ZapA^I83E^ localized throughout the cell, apparently unable to bind FtsZ at midcell (**Figures 2**, **S1**). FtsZ localized regularly at midcell for all cultures but also more diffusely for the Δ*zapA* and the ZapA^I83E^ cells. Since ZapB midcell localization is dependent on ZapA [7,14] and both proteins are reported to oscillate together between the cell poles [15], it was expected to localize diffusely through the cells expressing dimeric ZapA^I83E^. Instead ZapB localized polarly in these cells mimicking the pattern of cells without ZapA [7,15]. This sets ZapA^I83E^ apart from the other non-complementing ZapA mutants that did localize to midcell and recruited ZapB [7]. The ZapB localization pattern is a good indication whether ZapA complements at midcell (**Figure S2**).

**Figure 2.**
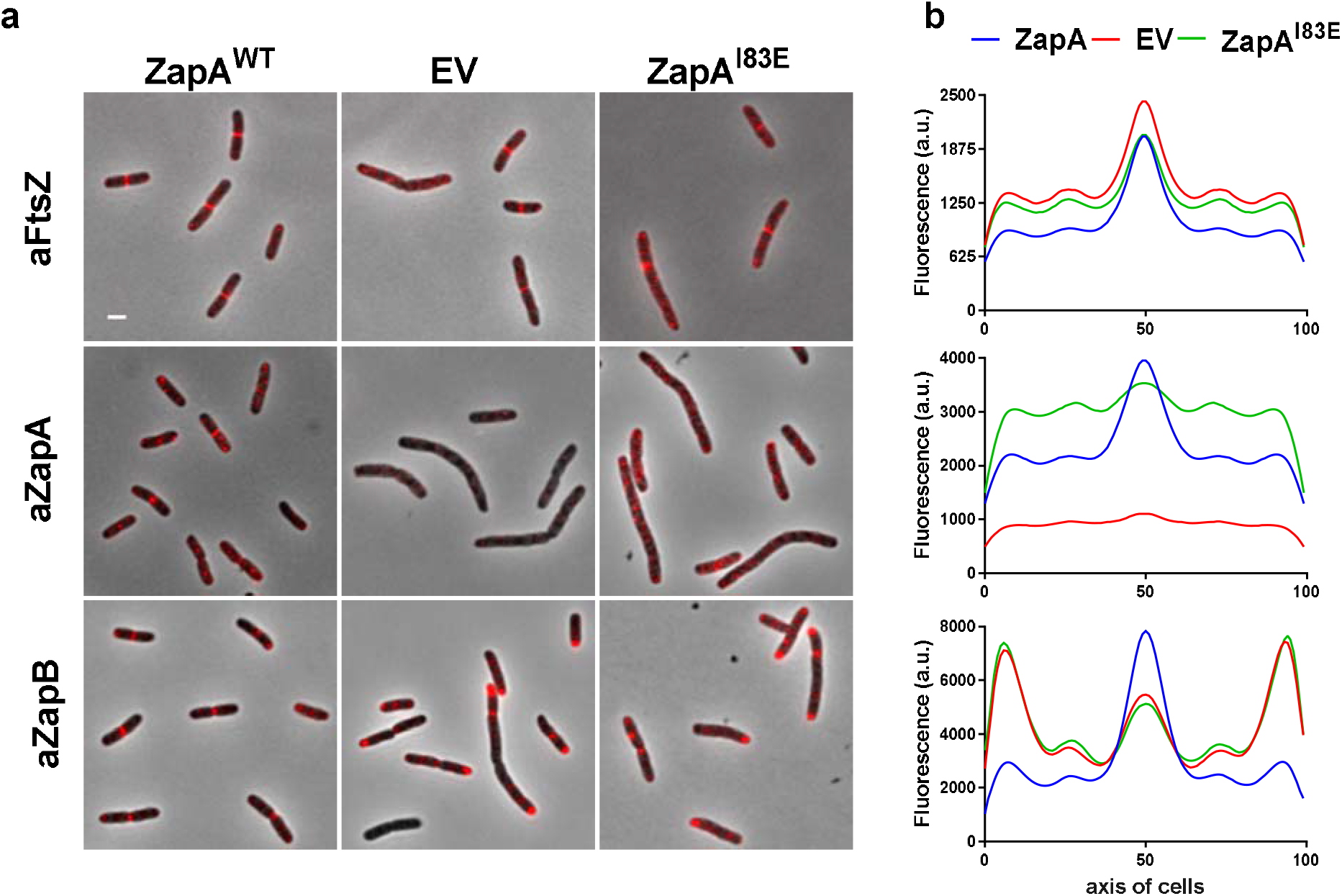
ZapA^I83E^ does not localize at midcell and does not complement based on ZapB localization. **(a)** TB28 Δ*zapA* cells expressing ZapA^WT^, an EV control or ZapA^I83E^ were immunolabeled with antiFtsZ, antiZapA and antiZapB and a fluorescent secondary antibody (red). The fluorescence signals from the cells were imaged and analyzed using ObjectJ [16]. The fluorescence intensities are shown at identical brightness and contrast values and the scale bar represents 2 μm. **(b)** The average fluorescence profiles along the cell axis reveal FtsZ midcell localization for all groups. ZapA^I83E^ is expressed and detectable and follows the localization distribution pattern of the EV negative control. Consequently, strong ZapB midcell localization was not observed for ZapA^I83E^ and EV. Maps of the fluorescence profiles are shown in **figure S1**.

### 2.3. Dimeric ZapA^I83E^ interacts with FtsZ in vivo

The *in vivo* ZapA^I83E^ results did not seem to confirm *in vitro* evidence that the dimer still binds FtsZ. However, its diffuse localization pattern does not exclude the possibility that it is still interacting with FtsZ. ZapA^I83E^ may interact with FtsZ that is therefore not capable to polymerize as a part of the Z-ring. Indeed, the lack of FtsZ bundling was one of the conclusions of the *in vitro* work using ZapA^I83E^ [6]. This should not necessarily have major effects on division given the large amount of FtsZ in the cell of which only 1/3 is involved in formation of the Z-ring. The critical concentration of FtsZ for polymerization *in vitro* is ~ 1 μM, whereas *in vivo* 5 μM is present. The amount of ZapA^I83E^ produced from plasmid is approximately that of the endogenous concentration of 0.4 μM as tetramer [7]. Not only is there 3 times as much FtsZ as ZapA in the cell, FtsZ overexpression if often observed as an attempt to compensate for division defects [17]. To assess the ZapA^I83E^-FtsZ interaction regardless of its localization pattern *in vivo*, a FRET experiment was attempted.

The interaction between mKO-FtsZ and mCh-ZapA has been demonstrated *in vivo* by FRET with 4.5 % energy transfer [18]. We aimed to compare the FtsZ-ZapA^WT^ interaction to the putative FtsZ-ZapA^I83E^ interaction. The FP-fusions to ZapA^WT^ and ZapA^I83E^ were confirmed to localize and complement exactly as described for the non-fused proteins under the conditions of a FRET experiment. In addition, the presence of endogenous ZapA did not change the localization pattern of ZapA^I83E^ suggesting that the mutant also cannot tetramerize in combination with ZapA^WT^ (**Supplementary text 5.1**, **Figure S3**). For the FRET experiment the Δ*zapA* strain was used to increase the chances of plasmid expressed mKO-FtsZ and mCh-ZapA to interact. The FRET experiment was performed as described [18,19]. The direct fusion between mKO and mCh forming a tandem as positive control gave 31 % energy transfer (E*f*A). The negative controls consisted of either mCh-ZapA, mCh-ZapA^I83E^ or mCh-PBP1b paired with the non-interacting IM protein mKO-PBP1a and gave low EfA values of less than 1.7 %. Together, these controls suggest a good detection range for the performed FRET experiments (**Figure 3**). Wild type mCh-ZapA interacted with mKO-FtsZ showing the expected energy transfer efficiency of 4 %. Interestingly, mCh-ZapA^I83E^ gave very similar EfA values and fluorescence spectra underscoring equal expression. The FRET experiments confirm the binding of dimeric ZapA^I83E^ to FtsZ previously shown *in vitro* by Pacheco-Gomes *et al* [6]. The FRET experiment was repeated in the WT strain resulting in similar EfA values of 4.5 and 5.5 % for ZapA and ZapA^I83E^, respectively.

**Figure 3.**
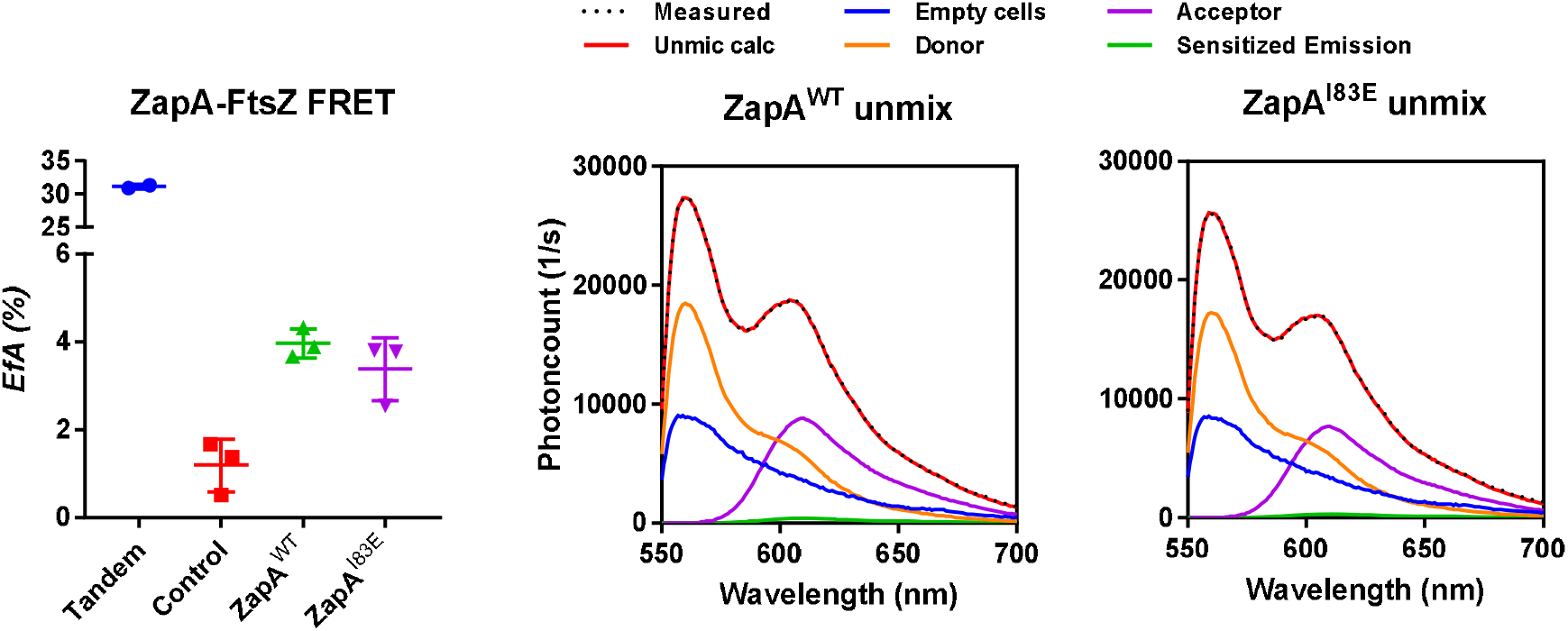
FtsZ and ZapA^I83E^ *in vivo* interaction FRET assay. On the left the energy transfer efficiency of the indicated FRET pairs. The positive and negative controls suggest a good experiment. On the right typical emission spectra plotted against the photocount for the mCh-ZapA and mKO-FtsZ pairs. The contribution of the direct excitation of mCh (purple) and of mKO (orange) and the background autofluorescence (blue) to the dotted measured spectrum are obtained by the unmixing procedure [18, 19]. The remaining signal (green) is the sensitized emission. mCh-ZapA^WT^ and mCh-ZapA^I83E^ showed similar fluorescence spectra with mKO-FtsZ resulting in very similar E*f*A values.

### 2.4. ZapB delocalizes unipolarly in cells with dimeric or absent ZapA

ZapB localizes at midcell in wild-type cells but polarly in the absence of ZapA [7,13,15]. In the presence of diffusely localized ZapA^I83E^, ZapB also localizes polarly instead of following the diffusely distributed pattern. To investigate, TB28 Δ*zapA* containing either ZapA^WT^, ZapA^I83E^ on plasmids or an EV negative control were grown under minimal medium conditions to prevent large differences in cell length (**Figure S4**). Expression was induced with 50 μM IPTG for at least 2 mass doublings and the cells were fixed and harvested for immunolabeling with antiZapB before imaging. The resulting fluorescence profiles showed the expected ZapB midcell localization in the cells expressing ZapA^WT^ and polar localization for cells expressing ZapA^I83E^ or EV (**Figure 4**). The ObjectJ “poleflipper” macro was used to orient the cell profiles based on the localization of the strongest ZapB signal, reorienting also the underlying channels [16]. This revealed still an average midcell localization of ZapB for the cells expressing ZapA^WT^ but also striking unipolar signals for the cells expressing ZapA^I83E^ or EV (**Figure 4b**). The polar localization of ZapB strongly suggest that ZapA tetramerization is a requirement for its interaction with ZapB.

**Figure 4.**
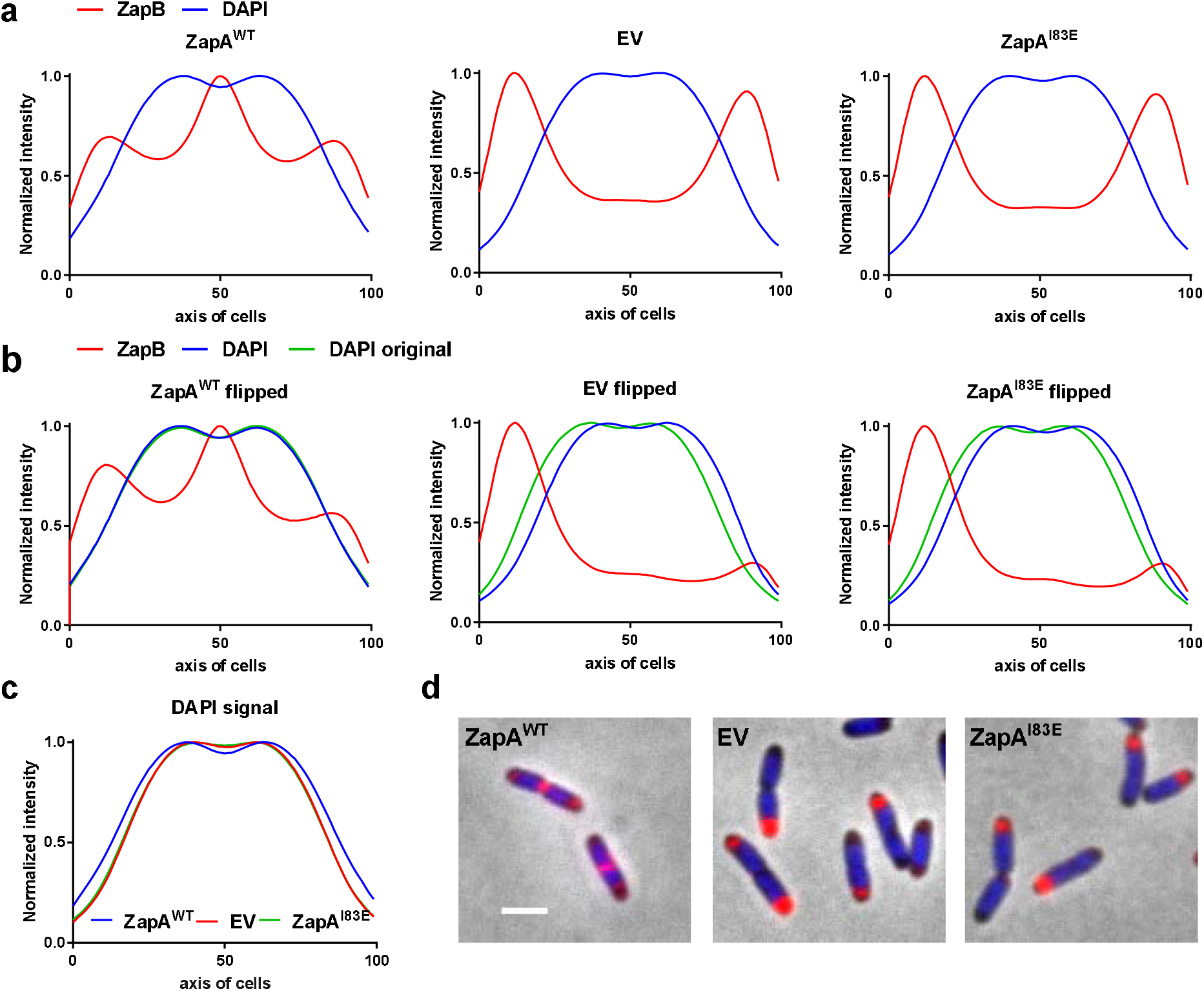
ZapB localizes unipolarly in Δ*zapA* or with plasmid expressed ZapA^I83E^ and is anti-correlating with the chromosomal position. **(a)** Localization profiles of immunolabeled ZapB and DAPI in Δ*zapA* expressing ZapA^WT^, EV or ZapA^I83E^. The ZapB profiles show strong signals at midcell in cells with functional ZapA and polar localization in cells without it or with dimeric ZapA. **(b)** When the cells are ordered based on strongest polar signal, predominantly unipolar signals are observed. The DAPI signal in these cells localizes away from ZapB. The blue and green DAPI profiles represent the flipped and non-flipped versions, respectively. **(c)** The DAPI signal in cells without ZapA or with dimeric ZapA seems to be more compacted towards the middle of the cells and shows less invagination. **(d)** Example of the ZapB immunolabeled (Red) cells with a DAPI stain (Blue). The scale bar represents 2 μm.

### 2.5 Unipolar ZapB signal anticorrelates with signal for the chromosome

ZapB interacts with MatP, which connects the chromosomal ter region to the divisome [10]. Possibly, its delocalization may influence the chromosome position. To verify this, the Δ*zapA* cells expressing ZapA^WT^, Zap^AI83E^ from plasmid or harboring an EV were visualized with DAPI (**Figure 4a**). When the ZapB poles are flipped, it becomes apparent that the chromosomal signal anti-correlates with the ZapB signal (**Figure 4b**). In the ZapB pole-flipped DAPI profiles for EV and ZapA^I83E^ the chromosome is localized away from ZapB while for cells with ZapA^WT^ the DAPI signal is unaffected. Overlaying the DAPI fluorescence profiles shows that the chromosomes are more localized towards midcell without strong nucleoid invagination for cells without ZapA or with dimeric ZapA^I83E^. Cells expressing ZapA^WT^ show more nucleoid invagination at the site where cell division requires proper nucleoid segregation (**Figure 4c**). The interaction partner of ZapB, MatP, indeed concentrates the ter region of the chromosome and a severed *ter* linkage may influence localization and compaction during cell division [11]. This may explain why the DAPI signals show less chromosome invagination in cells with uni-polar ZapB. These observations may provide further insights in ter linkage through ZapB function and chromosome segregation. The question remains whether the anti-correlated chromosome is a direct effect of polar ZapB or it is merely occupying the remaining space or do other actors direct the chromosome away from the pole.

### 2.6 Min oscillation is not affected by polar ZapB, polar ZapB is likely folded and functional

The multimerization state of ZapA is clearly important for *ter* linkage. Because dimeric ZapA is localizing diffusely in the cytoplasm, its oscillation seems to require its interaction with ZapB. However, ZapA seems not to be required for ZapB oscillation as it is still present in the poles and to a much lower extend at mid cell (as it is missing one of its binding partners) (**Figure S5**).

To proof that ZapB is not an inclusion body that is simply obstructing a pole, but is still dynamic, the Δ*zapA* strain and its parental TB28 wild type strain were immunolabelled with antiMinC. Min proteins oscillate from pole to pole where they inhibit Z-ring formation and by a concentration dependent manner allow the proto-ring to form at midcell. Previously, it was shown that the localization pattern of immunolabelled MinC reflects its oscillatory behavior [16]. The MinC pattern was identical in both strains, indicating that MinC completely ignored the presence of ZapB with respect to its position in the cell (**Figure 5**). These data suggest that the polar ZapB is properly folded and delocalized solely because its midcell recruiting partner ZapA is missing (or dimeric). Indeed, counter-oscillation of FtsZ, ZapA and ZapB has been proposed to build up the new cell division site influenced by Min [15].

**Figure 5.**
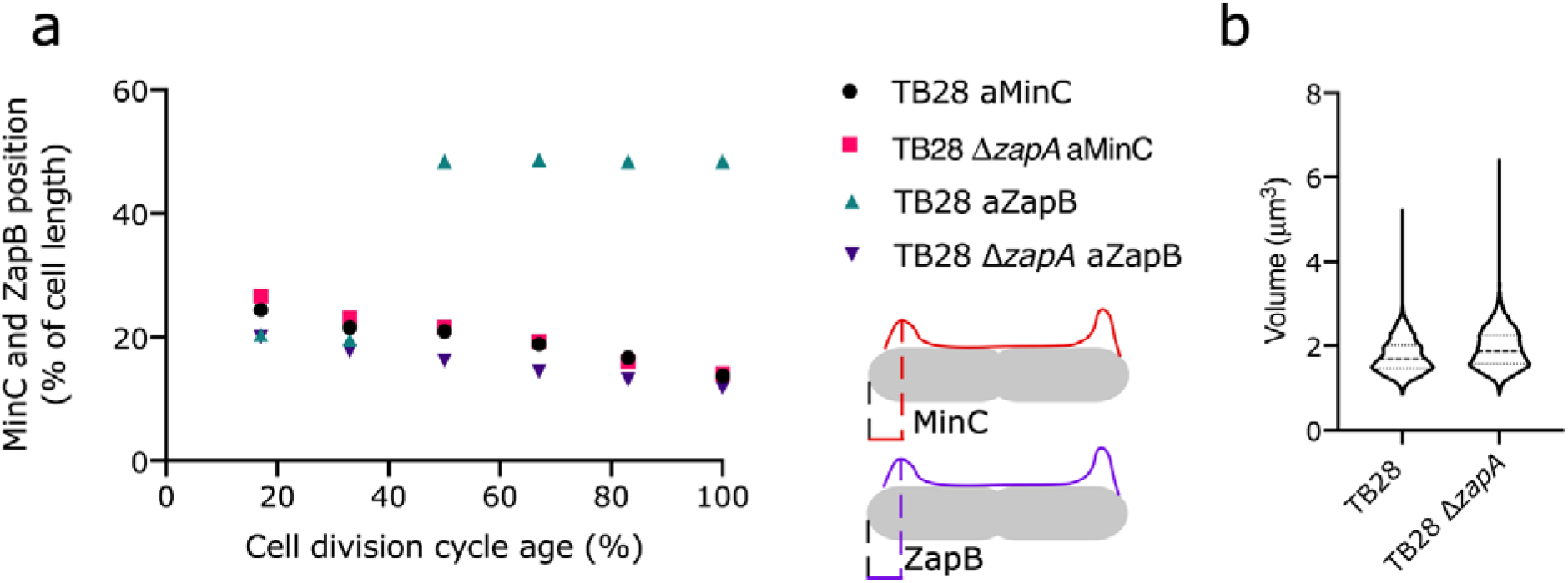
MinC localization is identical in TB28 and Δ*zapA* despite the polar presence of ZapB in the later strain. **(a)** The distance from the ZapB containing cell pole to the position of MinC in that cell pole was determined in TB28 Δ*zapA* and compared with the position of MinC in wild type cells and plotted against the normalized cell division cycle age (see also cartoon). The position of ZapB in the pole with the strongest ZapB signal in TB28 Δ*zapA* was also plotted and compared to its position in the wild type strain. In the wild type strain, all ZapB molecules end up at mid cell, whereas in the Δ*zapA* strain only a minor amount is present at mid cell (see for complete profiles **figure S5**). **(b)** Volume distribution of TB28 and TB28 Δ*zapA* cells. The number of cells analyzed were for TB28 aMinC, TB28 Δ*zapA* a(anti)MinC, TB28 aZapB, and TB28 Δ*zapA* aZapB 3201, 4154, 3473, and 1726, respectively.

### 2.7 ZapA and MatP effects on ZapB localization

Both ZapB and MatP are required to bring *matS* sites (and thus *ter*) to the division site [20]. If the physical interaction of ZapB and MatP is required for the correct localization of the chromosomes, changes in ZapB dynamics are expected in a Δ*matP* strain. Strain TB28 and Δ*zapA*, Δ*matP* or Δ*zapA* Δ*matP* in the isogenic background were grown in rich medium conditions before they were fixed, harvested and labeled with antiZapB. The fluorescence profiles were flipped and analyzed using ObjectJ (**Figure 6**). As expected, ZapB localized at midcell in the wild-type cells and mostly (uni)polarly in the Δ*zapA* cells. In the absence of MatP, ZapB was still able to be recruited to midcell and partly localized again in the poles and the cells’ periphery. The ZapB localization signals are substantially stronger in cells without MatP suggesting either more ZapB production by the cells or more ZapB epitope becomes available to antiZapB. The double deletion strain shows the combined effects of the single *zapA* or *matP* deletions. In these cells, ZapB localizes polarly but also more throughout the cell. This confirms that ZapA guides ZapB to midcell and suggests that MatP keeps ZapB from freely distributing throughout the cell.

**Figure 6.**
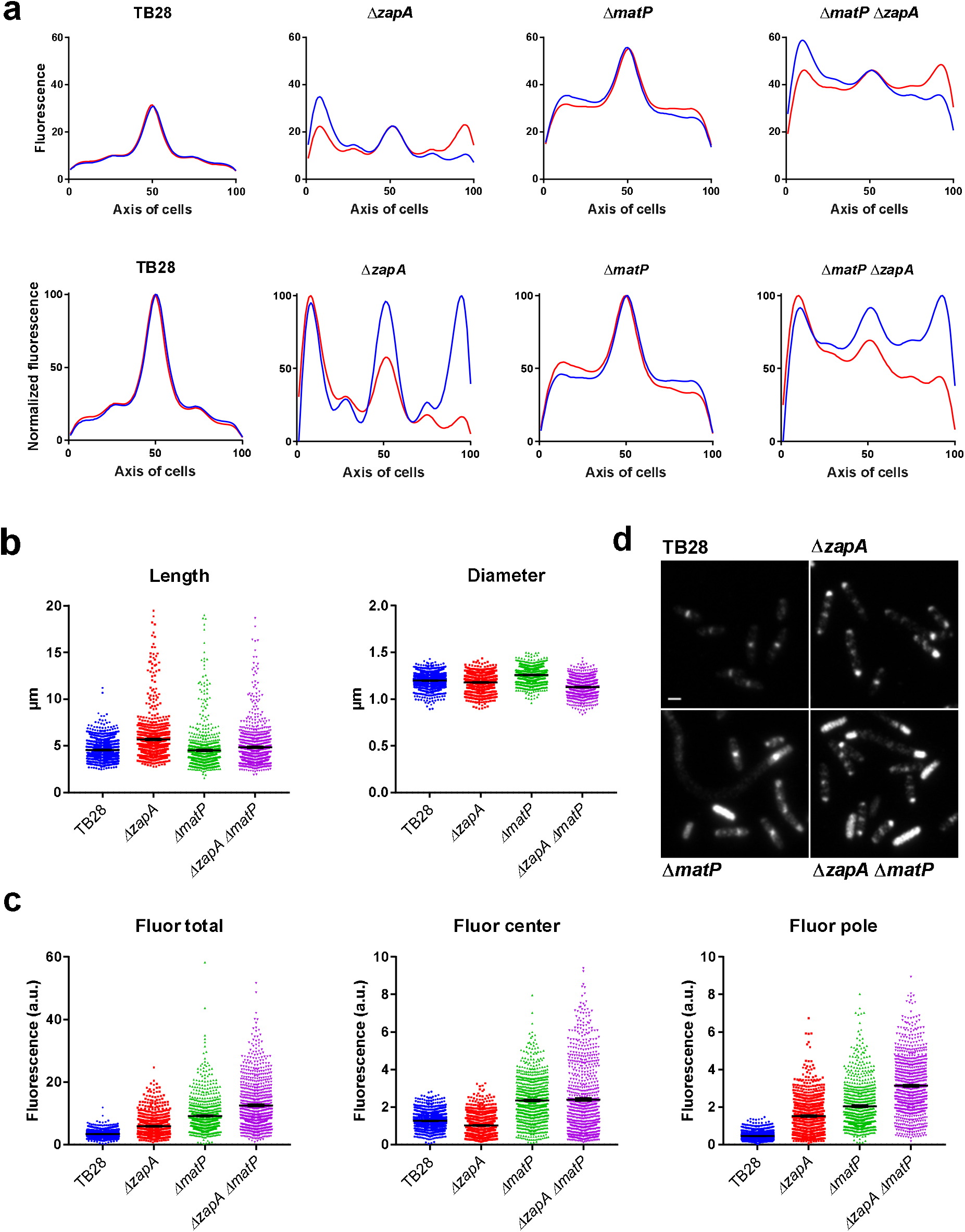
ZapA is the main factor affecting ZapB localization at midcell. **(a)** Immunolabeled ZapB profiles of TB28, Δ*zapA*, Δ*matP* or Δ*zapA* Δ*matP* grown in rich medium. Absence of MatP has minor effects in ZapB midcell localization but ZapB signals are stronger and localize more throughout the cell. **(b)** Analysis of cell length and diameter show the on average longer cells for Δ*zapA*. **(c)** Subcellular localization of ZapB shows delocalization patterns due to *zapA* and *matP* deletions. The error bars represent the SEM **(d)** Example of the immunolabeled cells. The fluorescence intensities are shown at identical brightness and contrast values and the scale bar represents 2 μm. The number of cells analyzed were for TB28, TB28 Δ*zapA*, TB28 Δ*matP*, and TB28 Δ*zapA* Δ*matP* 1471, 841, 746, and 820, respectively.

## 3. Discussion and conclusion

ZapA is an FtsZ associated protein involved in stabilization of the proto-ring and synchronizing bacterial cell division with chromosome segregation through *ter* linkage. ZapA can exist as tetramers or dimers and *in vitro* evidence suggests its functionality relies on its tetrameric form, even though the dimer can still bind to FtsZ [5,6]. Our work confirms this *in vivo* by investigating tetrameric and dimeric ZapA, as well as FtsZ at the cell division site, and ZapB as the link with chromosome segregation.

Although ZapA tetramerization is not a requirement for FtsZ binding, it is for FtsZ binding at midcell. The *in vivo* data suggest that dimeric ZapA only binds cytosolic FtsZ that may therefore be incapable of polymerizing into the proto-ring. Alternatively, the ZapA dimer may be unable to bind FtsZ filaments. The mutant to create dimeric ZapA (I83E), was designed to disrupt the C-terminal multimerization interface to yield parallel ZapA dimers with the globular head group intact but this exact conformation was not verified [6]. Parallel dimeric ZapA is expected to be capable of binding an FtsZ dimer and therefore associate with FtsZ in the proto-ring. Antiparallel ZapA dimerization would weaken the interaction with FtsZ dimers as well as with ZapB explaining the *in vivo* results presented here.

Tetrameric ZapA does not interfere with FtsZ protofilament formation and may even be required for its stability [6,12]. A remaining question is whether ZapA tetramerization itself or its interaction with FtsZ at midcell is required for ZapB midcell localization. Other ZapA mutants R13D, E51K and I56K were not complementing cell length but still localized at midcell as did ZapB [7]. This suggested they were interacting with FtsZ and independently still recruited ZapB, apparently affecting the proto-ring in a different manner. The recently published non-complementing ZapA^R46A^ mutation that has been shown to tetramerize but not bind FtsZ may help to answer this question [5,9].

ZapB becomes predominantly polar when ZapA is unable to interact with it. When its other interaction partner MatP is absent, more ZapB signal is detectable and still able to localize to midcell but more diffusely and in the cell poles. When both of ZapB’s interaction partners are missing, its localization pattern becomes more diffuse and bipolar. This suggests a role for MatP in the unipolar localization of ZapB in Δ*zapA* cells and continued dynamics if the *ter* linkage is broken. Although speculative, polar ZapB may have a function in WT cells at the beginning of the cell cycle. In this case, ZapB is part of the division ring (i.e. not aggregating) and may repulse the DNA after division. The negative charge of ZapB may be of influence on chromosome localization considering it high abundance in the cell [17] with its 17 negative and 8 positive residues this could be significant. However, presently no evidence exists for direct effects of ZapB on the chromosome. A potential way of re-recruitment to the new cell division site is through counter-oscialation with the other early cell division proteins. Only future experiments would lead to a better understanding of ZapB functionality and localization dynamics.

In conclusion, tetramerization of ZapA is required *in vivo* for its binding of FtsZ protofilaments in the Z-ring and for its interaction with ZapB.

## 4. Materials and Methods

### Bacterial strains and growth conditions

*Escherichia coli* K12 strains used in work are presented in **table 1**. The cells were cultured in rich medium (TY: 10 g Tryptone (Bacto laboratories, Australia), 5 g yeast extract (Duchefa, Amsterdam, The Netherlands) and 5 g NaCl (Merck, Kenilworth, NJ) per liter) supplemented with 0.5% glucose (Merck) or in glucose minimal medium (Gb1: 6.33 g K_2_HPO_4_ (Merck), 2.95 g KH_2_PO_4_ (Riedel de Haen, Seelze, Germany), 1.05 g (NH_4_)_2_SO_4_ (Sigma, St. Louis, MO), 0.10 g MgSO_4_·7H_2_O (Roth, Karlsruhe, Germany), 0.28 mg FeSO_4_·7H_2_O (Sigma), 7.1 mg Ca(NO_3_)_2_·4H_2_O (Sigma), 4 mg thiamine (Sigma), and 4 g glucose per liter, pH 7.0) at 28 °C while shaking at 205 rpm. For growth in Gb1 of TB28 based strains 50 mg lysine, 50 mg arginine, 50 mg glutamine. 20 mg uracil, and 2 mg thymidine (all from Sigma), were added per liter. Expression of protein was induced with isopropyl β-D-1-thiogalactopyranoside (IPTG, Promega, Madison WI) as indicated. Plasmids were maintained in the strains by addition of 100 μg.ml^−1^ ampicillin (Sigma) or 25 μg.ml^−1^ chloramphenicol (Sigma). Growth was measured by absorbance at 600 or 450 nm with a Biochrom Libra S70 spectrophotometer (Harvard Biosciences, Holliston, MA) for TY or Gb1 cultures, respectively. TB28 Δ*matP* and TB28 Δ*zapA* Δ*matP* were created by P1 transduction of *matp::kan* from the KEIO collection [21] into TB28 and TB28 Δ*zapA*, respectively. The presence of the deletions was confirmed by PCR.

**Table 1.**
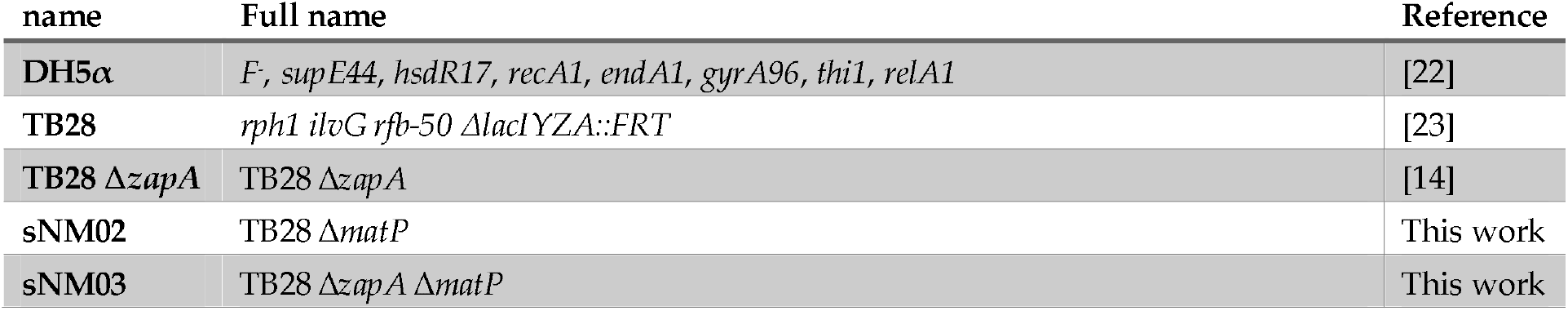
strains used

### Site-Directed Mutagenesis and Plasmid Construction

The plasmids used in this study are shown in **table 2**. Plasmid pGP021 [3] expresses ZapA from a weakened *trc99A* promotor, which is IPTG inducible. The I83E mutation was introduced in the ZapA expressing plasmid pGP016 (6HisZapA) using the Quick change mutagenesis method (Stratagene, la Jolla, CA, USA). The used primers were 5’-GTATGGAACAGCGTGAACGGATGCTGCAGC-3’ and 5’-GCTGCAGCATCCGTTCACGCTGTTCCATAC-3’ and the mutagenesis resulted in plasmid pET302His6ZapA^I83E^. From this plasmid ZapA^I83E^ was cut using NcoI and HindIII and inserted in pTHV037 to yield plasmid pRP071. Plasmid mCh-ZapA^I83E^ pNM137 was created by exchanging *zapA* in pSAV077 with zapA^I83E^ from pRP071 by restriction digestion cloning using DraIII and HindIII.

**Table 2.**
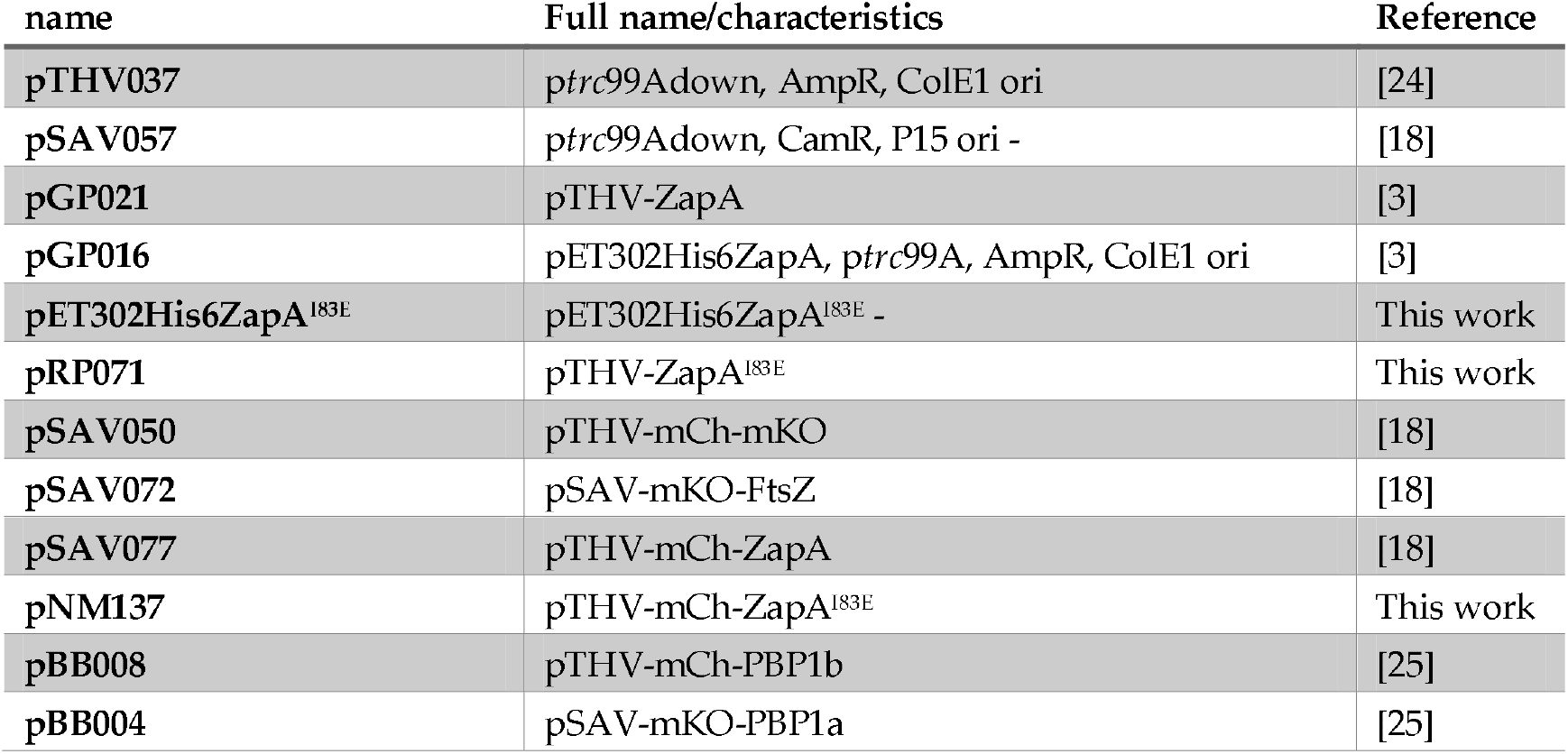
Plasmids used in this study

### Immunolabeling

Immunolabeling of cells was performed described [26] with antibodies against ZapA, FtsZ, or ZapB [16]. The ZapA antiserum was routinely purified by adsorption against TB28 Δ*zapA* cells to filter out potential cross-reactive IgG. The supernatant was subsequently used to label the ZapA mutants.

### Microscopy and Image analysis

For imaging, the cells were immobilized on 1 % agarose [27] and photographed with a CoolSnap fx (Photometrics) charge-coupled device (CCD) camera mounted on an Olympus BX-60 fluorescence microscope through an UPLANFl 100x/1.3 oil objective (Olympus,Tokyo, Japan). Images were taken using modified acquisition software that used the program ImageJ by Wayne Rasband and analyzed using Object-J’s Coli-Inspector [16].

### FRET assay

The FRET experiments were performed as originally described in [18,19] with the plasmids shown above.

## Supplementary Materials

Supplementary materials can be found at XXXX

## Author Contributions

Conceptualization, methodology, validation, formal analysis, investigation, data curation, writing—original draft preparation, writing—review and editing, visualization, supervision and project administration were by N.Y.M. and T.d.B.; funding acquisition by T.d.B.

## Acknowledgments

We are grateful to René van der Ploeg and Tjalling Siersma for creating the pTHV-ZapA^I83E^ expression plasmid and Elisa Galli for the antiZapB and Miguel Vicente for antiMinC. We thank Jolanda Verheul, Berivan Temiz and Eray Tarhan for technical assistance.

