## Supplementary Material for "ZapA tetramerization is required for midcell localization and ZapB interaction in *Escherichia coli*"

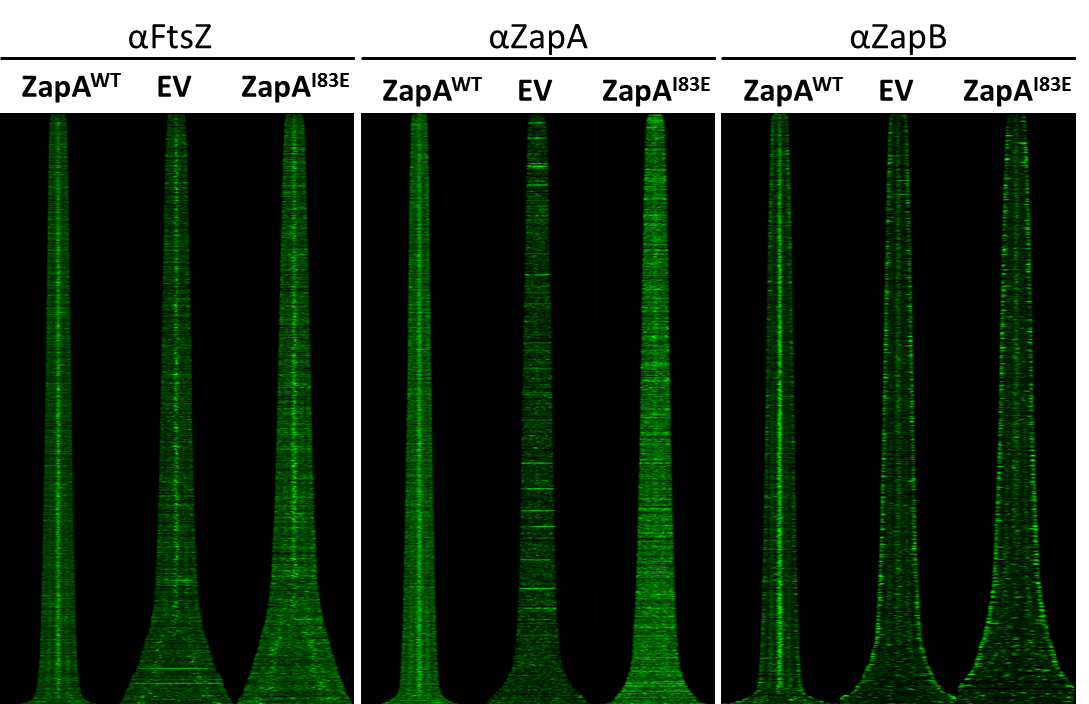

**Figure S1** – Map of fluorescence profiles of TB28 ∆*zapA* cells expressing WT ZapA, and empy vector control or ZapA^I83E^ immunolabeled with antibodies against FtsZ, ZapA or ZapB. Cells are sorted according to cell length. Average profiles are shown in the main text figure 2.

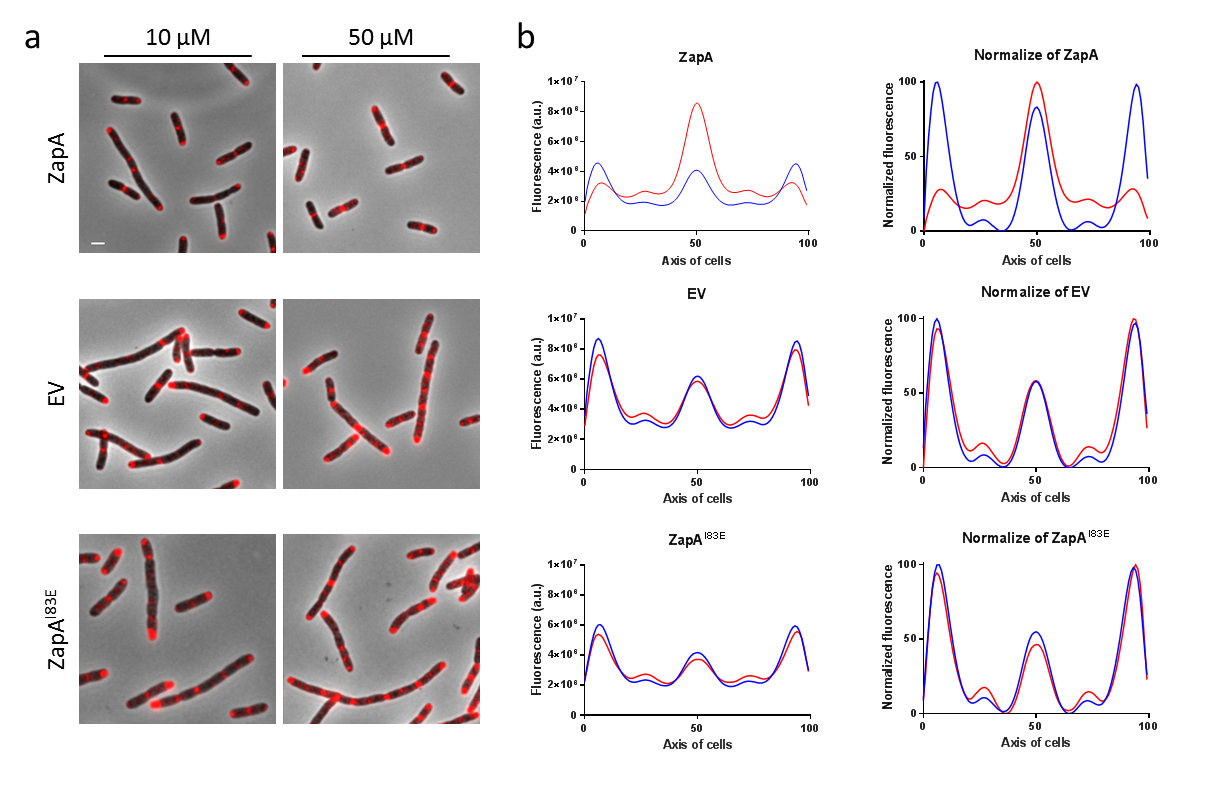

**Figure S2** – ZapA complementation can be shown by ZapB midcell localization. (**a**) At low induction concentrations ZapA function is not (fully) complemented, the cells become longer and ZapB localizes strongly in the cell poles even for the ∆*zapA* strain transformed with ZapA^WT^. When induced with 50 µM IPTG, ZapA^WT^ from plasmid is able to complement the delta phenotype resulting in normalized cell lengths and strong ZapB localization at midcell. The negative control of the ∆*zapA* strain transformed with EV does not complement the deletion phenotype at either induction concentration, the cells remain longer and ZapB localizes strongly in the cell poles. The non-tetramerizing ZapA mutant I83E is also unable to complement the ∆*zapA* phenotype and follows the same pattern as the EV cells suggesting it is non-functional in terms of cell length complementation and ZapB midcell localization. The scalebar represents 2 µm. (**b**) The average ZapB profiles of the 10 and 50 µM IPTG induced cells are shown by blue lines and red lines, respectively.

5.1. ZapA^WT^ and ZapA^I83E^ fusions do not further obstruct functionality

The mCh-ZapA^WT^ and mCh-ZapA^I83E^ constructs were tested for their use in a FRET experiment. First, their ability to complement the ∆*zapA* phenotype under minimal medium conditions was assessed. The mCh-ZapA^WT^ fusion complemented the cell length in ∆*zapA* strain with average cell lengths of 2.9 µm, which were very similar to the WT cells expressing the same construct. mCh-ZapA^I83E^ did not complement the *zapA* deletion strain and resembled the average cell length of the EV control cells more with 3.2 µm.

Localization patterns were observed in wild-type as well as in ∆*zapA* cells to ascertain whether chromosomal expression of ZapA would influence the localization of ZapA^I83E^. Additionally, this could indicate whether mCh-ZapA^I83E^ would be able to tetramerize with WT ZapA and potentially localize at midcell. mCh-ZapA^WT^ accumulates at midcell during cell division as is indicated by its increasing FCPlus (Surplus of fluorescence in cell center compared to the rest of the cell [1]) as a function of cell age, suggesting that it was functional (**Figure S3**). The mCh-ZapA^I83E^ mutant is more distributed throughout the cell. The fluorescence localization pattern of mCh-ZapA^I83E^ was identical in the WT and ∆*zapA* strain. This makes is it less likely that wild type ZapA and ZapA^I83E^ are able to form tetramers together.

mCh-ZapA^WT^ was able to recruit ZapB to midcell in WT as well as in ∆*zapA* cells underscoring its functionality albeit the ZapB signal was less pronounced. The latter observation may suggest that the FP fusion may be less favorable for ZapB binding. mCh-ZapA^I83E^ did not allow ZapB localization at midcell, completely mimicking the *zapA* deletion strain. ZapB localization in the WT strain was unhindered by the expression of ZapA^I83E^. This again suggests that ZapA^WT^ and ZapA^I83E^ do not form tetramers together. It was concluded that the constructs could be used for a FRET experiment assaying their interaction with FtsZ.

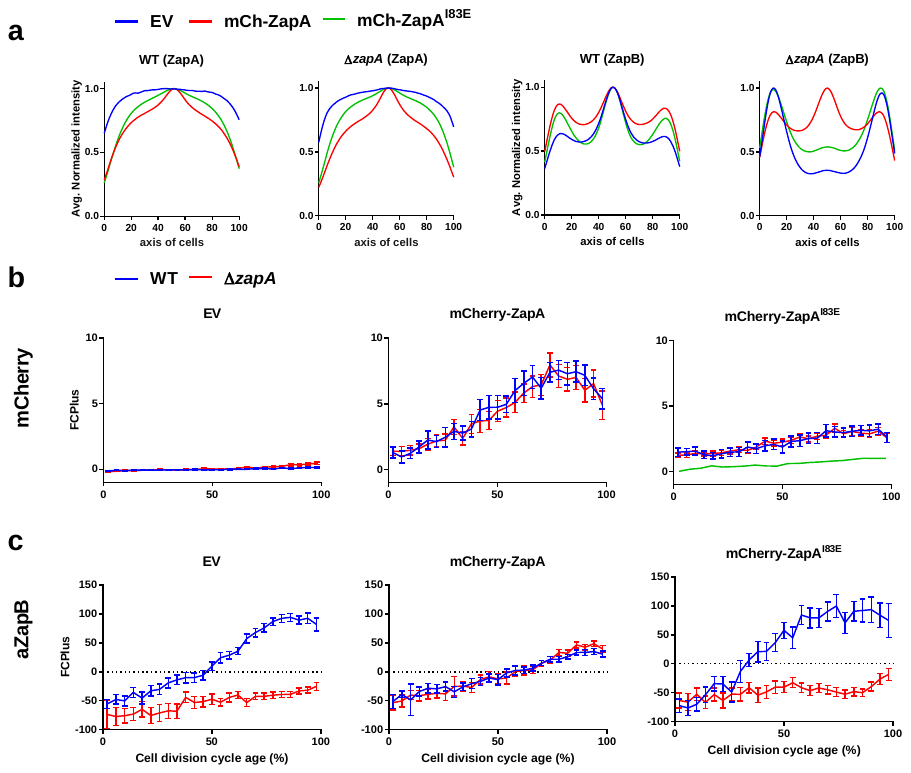
**Figure S3** – mCh-ZapA independently localizes to midcell and recruits ZapB but mCh-ZapA^I83E^ does not show strong affinity for midcell and has lost the ability to recruit ZapB. (**a**) Localization patterns of mCh-ZapA and mCh- ZapA^I83E^ in WT and ∆*zapA* cells and their ZapB immunolabeling. (**b**) mCh FCPlus values of steady state growing TB28 and TB28 ∆*zapA* expressing mCh-ZapA, mCh-ZapA^I83E^ or carrying the EV control. In both the WT and ∆*zapA* strain mCh-ZapA midcell concentrations increase as the cell division cycle progresses reaching the maximum concentration at around 70-80%. mCh-ZapA^I83E^ shows a less pronounced midcell localization similar to randomly diffused mCherry (green line). (**c**) The same cells as for the mCh measurement were immunolabeled with antiZapB showing the WT-ZapB response on mCh-ZapA or the mutant. In the WT cells ZapB localizes maximally at midcell around 80% of the cell cycle while in the ∆*zapA* strain it keeps a polar fraction and is not able to bind ZapA at midcell anymore with the resulting negative FCPlus values. Expression of mCh-ZapA enables ZapB to localize at midcell but mCh-ZapA^I83E^ does not. FCPlus values are plotted along the cell division age and binned into 4 % age groups. The bar represents the 95% confidence interval. Between 1000 and 5000 cells were analyzed for each group.

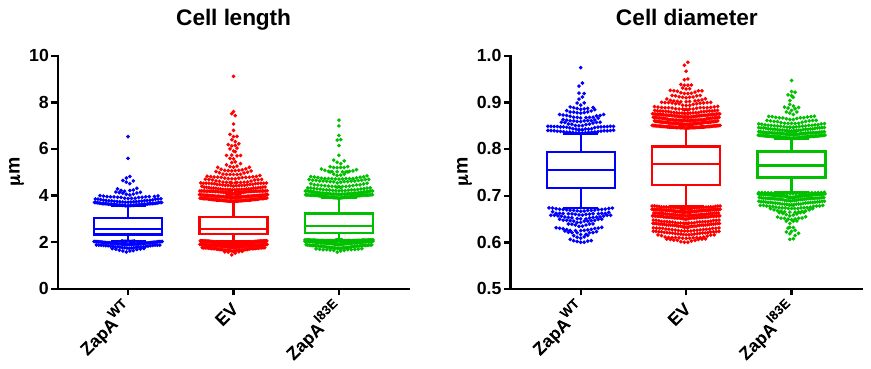

**Figure S4** – Cell lengths and diameters of the ZapB and DAPI labeled cells presented in **figure 4**. TB28 ∆*zapA* cells were grown at semi steady state in Gb1 at 28 °C and expression of ZapA^WT^, ZapA^I83E^ or EV (empty vector control) were induced for at least two mass doublings with 50 µM IPTG. The cells were fixed and immunolabeled with antibodies against ZapB and a fluorescent secondary antibody. The chromosomes were visualized by DAPI staining. The average cell lengths (in µm) were for ZapA^WT^ 2.72, EV 2.79 and ZapA^I83E^=2.88 and the respective number of measured cells were 1089, 3213 and 3213. The whiskers represent the 10^th^ and 90^th^ percentiles.

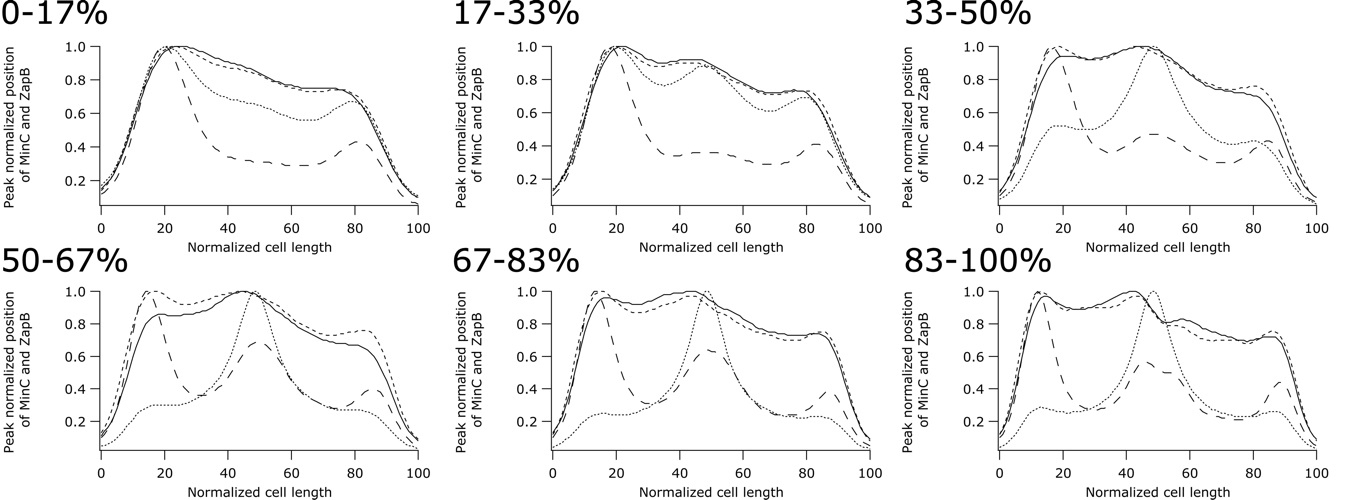

**Figure S5** - MinC is unaffected by the presence of ZapB with respect to is oscillation position in the Δ*zapA* cells. Peak normalized binned average profiles of ZapB and MinC in the indicated age classes plotted against the normalized cell length. The cells were grown in AB medium supplemented with glucose as carbon source [2]. Solid lined TB28 antiMinC, dashed lines Δ*zapA* antiMinC, dotted lines TB28 antiZapB and larger dash lines Δ*zapA* antiZapB,
